# Transcriptomic profiling of long- and short-lived mutant mice implicates mitochondrial metabolism in ageing and shows signatures of normal ageing in progeroid mice

**DOI:** 10.1101/2020.10.20.347013

**Authors:** Matias Fuentealba, Daniel K. Fabian, Handan Melike Dönertaş, Janet M. Thornton, Linda Partridge

**Affiliations:** Institute of Healthy Ageing, Department of Genetics, Evolution and Environment, University College London, London WC1E 6BT, UK; European Molecular Biology Laboratory, European Bioinformatics Institute, Wellcome Genome Campus, Hinxton CB10 1SD, UK; Max Planck Institute for Biology of Ageing, Cologne, Germany

**Keywords:** ageing, gene expression, lifespan, mouse, progeria, mitochondria, metabolism

## Abstract

Genetically modified mouse models of ageing are the living proof that lifespan and healthspan can be lengthened or shortened, yet the molecular mechanisms behind these opposite phenotypes remain largely unknown. In this study, we analysed and compared gene expression data from 10 long-lived and 8 short-lived mouse models of ageing. Transcriptome-wide correlation analysis revealed that mutations with equivalent effects on lifespan induce more similar transcriptomic changes, especially if they target the same pathway. Using functional enrichment analysis, we identified 58 gene sets with consistent changes in long- and short-lived mice, 55 of which were up-regulated in long-lived mice and down-regulated in short-lived mice. Half of these sets represented genes involved in energy and lipid metabolism, among which *Ppargc1a*, *Mif*, *Aldh5a1* and *Idh1* were frequently observed. Based on the gene sets with consistent changes and also the whole transcriptome, we observed that the gene expression changes during normal ageing resembled the transcriptome of short-lived models, suggesting that accelerated ageing models reproduce partially the molecular changes of ageing. Finally, we identified new genetic interventions that may ameliorate ageing, by comparing the transcriptomes of 51 mouse mutants not previously associated with ageing to expression signatures of long- and short-lived mice and ageing-related changes.

**Highlights:**

- Transcriptomic changes are more similar within mutant mice that show either lengthened or shortened lifespan
- The major transcriptomic differences between long- and short-lived mice are in genes controlling mitochondrial metabolism
- Gene expression changes in short-lived, progeroid, mutant mice resemble those seen during normal ageing

## 1. Introduction

Ageing is a complex phenotype. In humans, it causes a gradual decline in physiological function, and it is the major risk factor for many serious diseases in developed countries (MacNee et al., 2014; Niccoli and Partridge, 2012; Partridge et al., 2018). The rate of ageing is influenced by hundreds of genes, but it can be modulated in mice and other model organisms by single gene mutations (Bartke, 2011; Folgueras et al., 2018; Kenyon, 2005; Piper and Partridge, 2018; Uno and Nishida, 2016). Genetically modification of mice has been used to target each of the hallmarks of ageing (Folgueras et al., 2018; López-Otín et al., 2013). For example, mutations in genes encoding components of the nutrient sensing pathways can generate mice that outlive their wild-type littermates and display lower incidence of age-related diseases (Brown-Borg et al., 1996; Coschigano et al., 2000; De Magalhaes Filho et al., 2017; Flurkey et al., 2001; Hofmann et al., 2015; Holzenberger et al., 2003; Selman et al., 2009; Sun et al., 2013). In contrast, mutant mouse models of increased genomic instability have been suggested to be characterised by accelerated ageing, with shortened lifespan and early onset of age-associated pathologies (Dollé et al., 2011; Osorio et al., 2011; Van Der Pluijm et al., 2007; Weeda et al., 1997).

Transcriptomic studies have been widely used to identify molecular mechanisms associated with mammalian ageing, for example by comparing different species (eg. Fushan et al. 2015, Yu et al. 2011) or mouse strains (Houtkooper et al., 2013; Swindell, 2007) with different lifespans. Additionally, several studies in mice have revealed differentially expressed genes and pathways in response to genetic (Boylston et al., 2006; Rowland et al., 2005; Selman et al., 2009; Zhang et al., 2012), pharmacological (Fok et al., 2014b; Martin-Montalvo et al., 2013) and dietary (Rusli et al., 2015; Swindell, 2009) interventions that affect ageing. The transcriptomes of pairs of lifespan-extending interventions have also been compared, for instance, treatment with rapamycin and caloric restriction (CR) (Fok et al., 2014a), long-lived Ames and Little dwarf mice (Amador-Noguez et al., 2004) and CR and Ames dwarf mice (Tsuchiya et al., 2004), revealing conserved changes in gene expression. Comparison of multiple lifespan-extending interventions, including Snell, Ames, and Little dwarf mice together with CR and CR-mimetic compounds have shown that gene signatures associated with increased lifespan are enriched for steroid metabolism, cell proliferation, and cellular morphogenesis (Swindell 2007) or oxidative phosphorylation, drug metabolism and immune response (Tyshkovskiy et al., 2019). Comparison of mouse models with accelerated ageing revealed conserved gene expression changes in genes involved in lipid and carbohydrate metabolism, cytochrome P450s and developmental regulation of transcription (Kamileri et al., 2012).

A further approach to pinpoint the molecular mechanisms that are causal for changes in lifespan is to identify genes and pathways whose expression changes in opposite directions in long- and short-lived mouse strains. Paradoxically, one study making such comparisons found that long- and short-lived mice share a reduction in growth hormone/insulin-like growth factor 1 signalling and an increase in antioxidant responses, with no major transcriptomic differences (Schumacher et al., 2008). Since that study, further transcriptomic studies of mouse models of ageing have been published, providing the opportunity for further comparison of the changes in gene expression in long- and short-lived strains.

An additional layer of information to identify molecular mechanisms associated with lifespan is the changes in gene expression that occur during ageing. Several studies have been made of various mouse tissues (reviewed in Stegeman & Weake 2017; Frenk & Houseley 2018). For instance, Jonker et al. 2013 analysed 5 tissues (liver, kidney, spleen, lung and brain) at 6 time points (13, 26, 52, 78, 104 and 130 weeks) and found that, besides expression changes in immune response genes, most transcriptomic changes are organ-specific and involve alterations in the cell cycle, DNA repair, energy homeostasis and reactive oxygen species. More recently, single-cell RNA sequencing of kidney, lung and spleen mouse cells across age revealed a conserved down-regulation of protein translocation and up-regulation of antigen processing and inflammatory pathways (Kimmel et al., 2019).

In recent years, several *in silico* methods have been developed for the prediction of genetic interventions to delay ageing (reviewed in Fabris et al. 2017). For example, Huang et al. (2012) used protein interaction networks and machine learning approaches to predict the effect of gene deletions on yeast lifespan. Wan et al. (2015) used gene ontology terms comprising ageing-related genes to classify *Caenorhabditis elegans* genes into anti- or pro-longevity. We have taken a slightly different approach, by comparing the transcriptomes of long- and short-lived mice against gene expression data from genetic interventions not previously associated with ageing.

Here, we analysed and compared transcriptomic data from long- and short-lived mutant mice to identify molecular mechanisms associated with lengthening and shortening of lifespan. By further comparing the transcriptomes of mouse models of ageing to the transcriptomic changes observed during normal ageing, we determined which interventions resemble or reverse normal ageing-related changes. Finally, we identified novel genetic interventions with the potential to ameliorate ageing, by comparing all transcriptomic data publicly available from genetic interventions in mice against the changes in gene expression observed in mice with delayed, accelerated and normal ageing.

## 2. Results and discussion

### 2.1. Similarities and differences in patterns of transcriptional changes in long- and short-lived mutant mice

We first asked if genetic interventions that lengthen or shorten lifespan show characteristic transcriptomic changes, by comparing publicly available microarray and RNA-seq data from 10 long-lived and 8 short-lived mouse models of ageing **(Figure 1)**. The data came from 26 independent studies and included samples from adipose, brain, liver and muscle. To avoid potential batch effects, we derived 57 independent datasets using each study, genotype, tissue, sex and age, and performed differential expression analysis **(Supplementary Table 1)**. We then calculated the Spearman’s rank correlation coefficient (*r_s_*) between the gene expression fold changes **(Supplementary Figure 1)**. Given that the transcriptomes from several mutants were measured in multiple studies and within each study in multiple datasets, we averaged the correlations for the same genetic interventions **(see Methods 4.3)**. Since thousands of genes were used to calculate these correlations, even small correlations were statistically significant. To better estimate a threshold of significance for transcriptome-wide correlations, we analysed and correlated 65 transcriptomic datasets covering 51 genetic interventions in mice not previously associated with ageing **(Supplementary Table 2)**. We observed that 5% of the comparisons between the various genetic interventions had an absolute correlation coefficient higher than 0.15 **(Supplementary Figure 2)**. Thus, transcriptome-wide correlations above this value were considered statistically significant for the mutants affecting lifespan **(black squares in Figures 2A, C-E)**.

**Figure 1.**
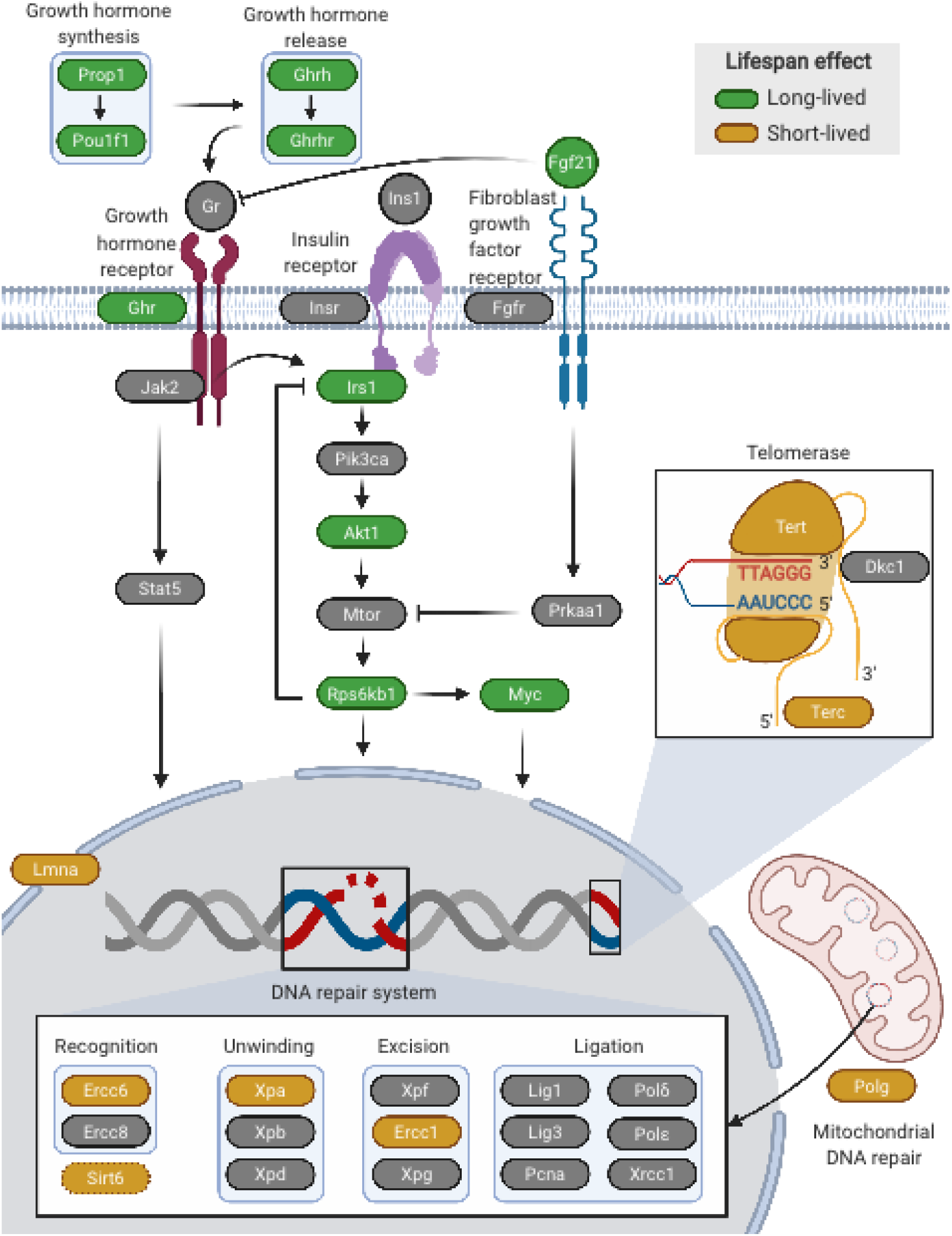
Pathways and processes modulated by the mouse models of ageing with transcriptomic data available. Genes whose mutation lengthen or shorten lifespan are coloured in green and yellow, respectively. Figure created with BioRender.com.

**Figure 2.**
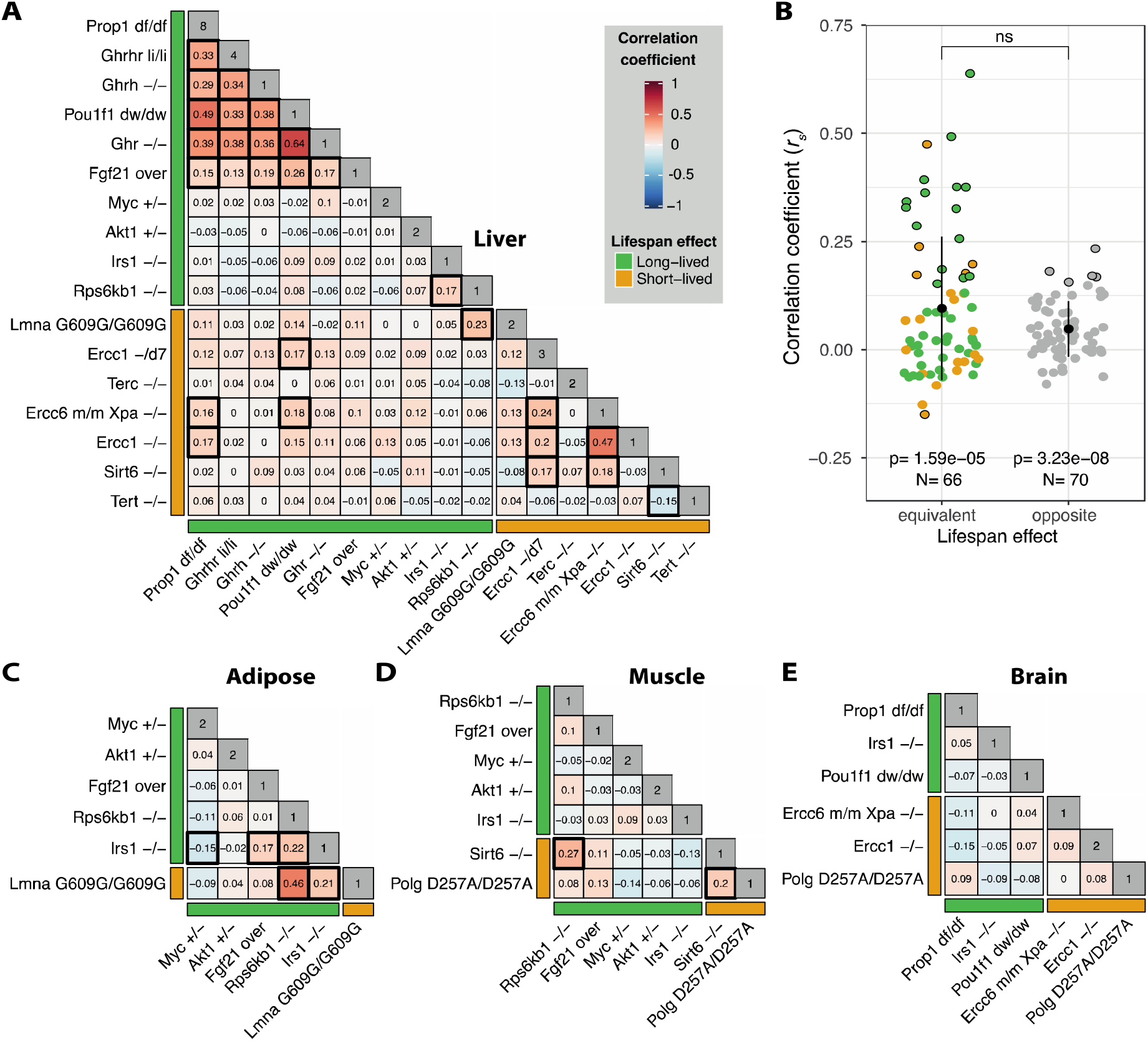
(A) Spearman’s rank correlation coefficients between the liver transcriptome of the mouse models of ageing. The intensity of the colours represents the magnitude of the correlations. Bars adjacent to the heatmaps indicate the effect on lifespan from each intervention. Diagonals show the number of datasets associated with each intervention. Black squares mark statistically significant correlations (i.e. |*r_s_*| > 0.15). (B) Pairwise correlations between the liver transcriptomes of interventions with equivalent or opposite impact on lifespan. Error bars represent one standard deviation from the mean. P-values below were calculated using a t-test with a population mean equal to zero as the null hypothesis. Statistical significance at the top is for the difference between the groups calculated using unpaired, two-samples Wilcoxon-test. Points circled in black represent statistically significant correlations. Transcriptome correlations between mouse models of ageing in the (C) adipose, (D) muscle and (E) brain. Heatmaps follow the same scheme used in panel A.

In the liver, 65% of correlations between interventions with equivalent effects on lifespan (i.e. long-lived vs long-lived and short-lived vs short-lived), were positive, reaching similarities as high as *r_s_* = 0.64 (*Ghr^−/−^* vs *Pou1f1^dw/dw^*) within long-lived mice and *r_s_* = 0.47 (*Ercc6^m/m^Xpa^−/−^* vs *Ercc1^−/−^*) within short-lived mice **(Figure 2A)**. In long-lived mice, mutations in genes controlling the synthesis and release of growth hormone and the growth hormone receptor **(Figure 1, top left)** induced remarkably similar transcriptomic changes (mean *r_s_* = 0.33). *Fgf21* over-expression also induced a similar transcriptome to the growth-hormone-related mutants, probably due to its well-known role in inhibiting growth hormone signalling (Inagaki et al., 2008). There were also positive correlations between interventions in the insulin signalling pathway, particularly between *Irs1^−/−^* and *Rps6kb1^−/−^* mutants, which is interesting considering that *Rps6kb1* directly phosphorylates and inhibits *Irs1* (Zhang et al., 2008). Although it is well known that growth hormone activates the insulin signalling pathway via *Irs1*, we did not detect any significant correlation between *Irs1^−/−^* and the growth-hormone-related mutants, possibly because of effects of growth hormone signalling on additional downstream targets to insulin signalling. Heterozygous mutation of *Akt1*, which is also involved in insulin signalling, did not induce a similar expression pattern to other long-lived mice, possibly reflecting the pleiotropic effects of *Akt1* function and hence alternative mechanisms to prolong lifespan. Among short-lived models, the highest correlation was observed between interventions in the DNA repair system **(Figure 1, bottom)**, including *Ercc6^m/m^Xpa^−/−^*, *Ercc1^−/−^* and *Ercc1^−/d7^* (mean *r_s_* = 0.24). *Sirt6^−/−^* also showed similarity with *Ercc1^−/d7^* and *Ercc6^m/m^Xpa^−/−^*, which may be explained by the recently discovered function of *Sirt6* as a DNA strand break sensor and activator of the DNA damage response (Onn et al., 2020). The only statistically significant negative correlation we found was between *Tert* and *Sirt6* knockout mice. With some exceptions, these results suggest that mutations in genes whose product participate in the same process or directly interact within the same signalling pathway are more likely to induce similar transcriptomic changes.

Surprisingly 78% of the correlations between interventions leading to opposite effects on lifespan (i.e. long-lived vs short-lived) were also positive, but smaller than interventions with similar lifespan effects. The maximum correlation observed of this type was an unreported one between *Lmna^G609/G609G^* and *Rps6kb1^−/−^* (*r_s_* = 0.23). We also observed a correlation between interventions in the DNA repair system (i.e. *Ercc6^m/m^Xpa^−/−^*, *Ercc1^−/−^* and *Ercc1^−/d7^*) and mutants of genes regulating the synthesis of growth hormone (i.e. *Pou1f1^dw/dw^* and *Prop1^df/df^*), which has been previously attributed to similar gene expression patterns on the somatotropic axis and anti-oxidant responses (Schumacher et al., 2008). In summary, although many correlations were positive even when the effects on lifespan were opposite, correlations between interventions with equivalent effects on lifespan were more likely to reach the threshold of significance **(Figure 2B)**.

We next assessed if the correlations we observed in the liver were also present in other tissues. From the 51 pairwise correlations in the other tissues, 22 (43%) followed the same direction as in the liver, but 18 (35%) were in the opposite direction. The remaining 11 pairwise correlations corresponded to comparisons with the mutant *Polg^D257A/D257A^*, for which there was no data available from the liver. Among the positive and significant correlations observed in the liver (i.e. *r_s_* > 0.15) only the comparisons between *Lmna^G609G/G609G^* vs *Rps6kb1^−/−^* and *Irs1^−/−^* vs *Rps6kb1^−/−^* were significant in other tissues (**Figure 2C**, *r_s_* = 0.46 and 0.22, respectively). The only negatively correlated interventions in the liver involved *Tert^−/−^*, which was not measured in other tissues. We also found tissue-specific correlations that were not significant in the liver, including *Rps6kb1^−/−^* vs *Sirt6^−/−^* in the muscle (*r_s_* = 0.27) **(Figure 2D)** and *Irs1^−/−^* vs *Lmna^G609G/G609G^* in the adipose tissue (*r_s_* = 0.21). Likewise, some correlations found in the liver were not detected in the brain such as *Pou1f1^dw/dw^* vs *Prop1^df/df^* and *Ercc1^−/−^* vs *Ercc6^m/m^Xpa^−/−^* **(Figure 2E)**. This analysis indicates that interventions into ageing may induce tissue-specific effects.

### 2.2. Functional analysis of transcriptional changes in mutant mouse models of ageing

We next sought to identify pathways enriched in the gene expression changes observed in the mutant mouse models of ageing. We only employed the liver data for this analysis, as it was the only tissue with enough interventions to identify statistically significant trends. We performed functional enrichment analysis on each dataset using Gene Ontology (GO) terms and a Gene Set Enrichment Analysis (GSEA). To identify gene sets (i.e. GO terms) with consist changes, we calculated their median rank based on enrichment scores across interventions and then we compared it against a random distribution of median ranks from the same number of interventions **(see Methods 4.4)**.

After multiple testing correction, we identified 470 gene sets in long-lived mice **(Supplementary Table 3)**, and 99 gene sets in short-lived mice showing consistent changes **(Supplementary Table 4)**. Remarkably, 93% of the gene sets found in short-lived mice were down-regulated, whereas 57% of the gene sets identified in long-lived mice were up-regulated. Surprisingly, 58 gene sets showed consistent changes in both groups of mice **(Figure 3A)**, of which 55 were up-regulated in long-lived mice and down-regulated in short-lived mice. These gene sets included 36 biological processes, 10 of which were linked with energy metabolism and 6 with lipid metabolism. We also observed the same trends in several processes associated with the metabolism of drugs, nucleic acids, amino acids and carboxylic acid. Consistent with the alteration in energy metabolism, we found similar gene expression patterns in genes forming the mitochondrial membrane and the electron transport chain, as well as genes coding for proteins with NADH dehydrogenase activity. Overall, these transcriptomic changes match well with previous studies in long-lived mice reporting an increase in protein levels and activity of several components of the electron transport system, as well as increased physiological markers of mitochondrial metabolism (Anderson et al., 2009; Brown-Borg et al., 2012; Westbrook et al., 2009). Also, the increased expression of genes controlling lipid metabolism is biologically meaningful, considering that previous studies have found that long-lived mice tend to use fat as an energy source, instead of carbohydrates (Westbrook et al., 2009).

**Figure 3.**
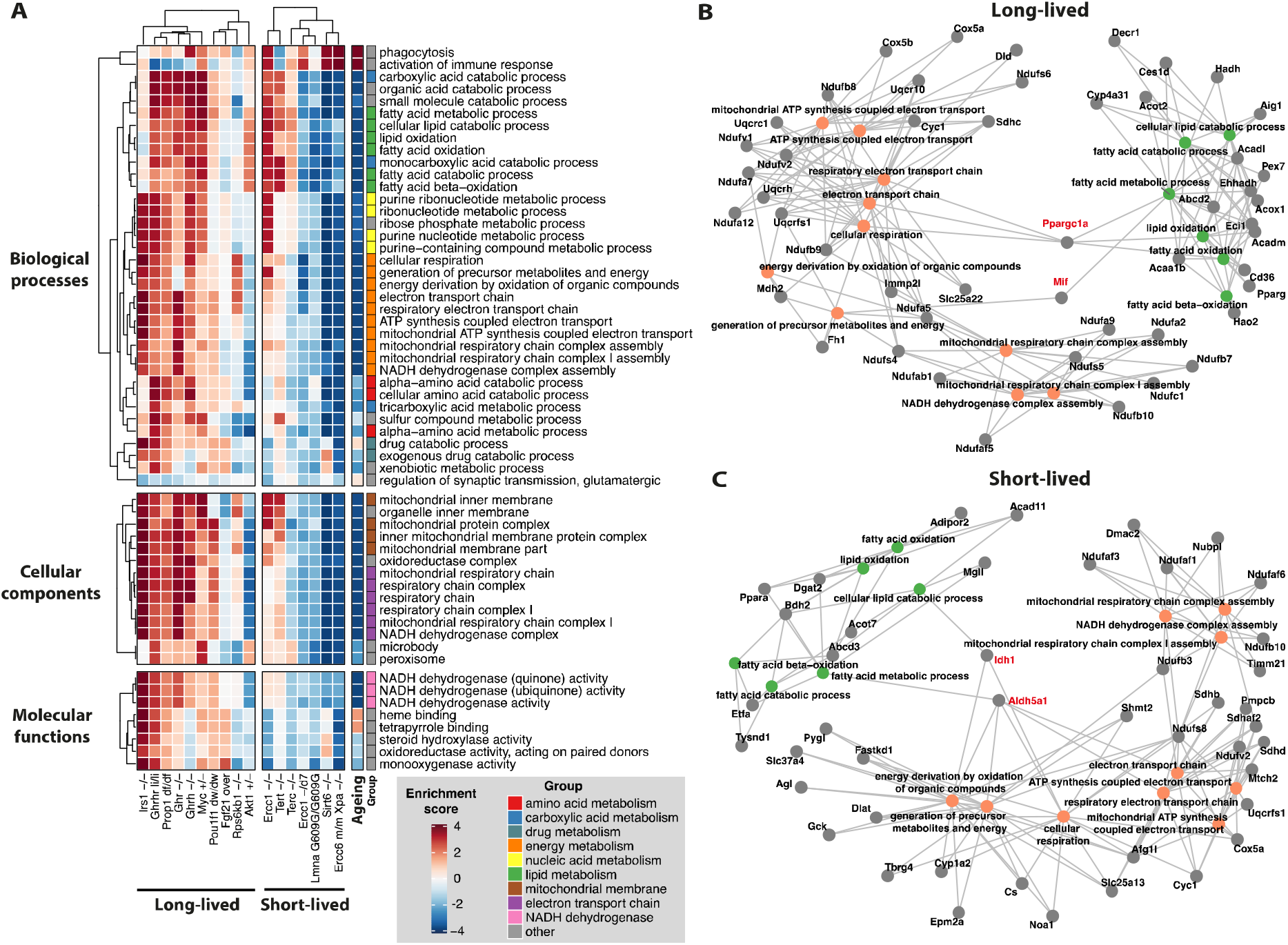
(A) Gene sets showing consistent changes in long- and short-lived mice (FDR < 0.05). The heatmap colours represent the statistical significance of the enrichment and the direction of the change. The ‘Ageing’ column represents the enrichment scores associated with the transcriptomic changes during ageing. The ‘Group’ column indicates different groups of gene sets with similar function. Genes involved in the regulation of biological processes linked with energy and lipid metabolism in (B) long-lived and (C) short-lived mice. Labelled in red are genes involved in both sets of processes. Nodes representing genes are coloured in grey, and nodes from biological processes are coloured based on the groups in panel A.

Based on a leading-edge analysis, we asked which genes contributed the most to the changes in gene expression observed in energy and lipid metabolism and if some genes were acting as hubs between both sets of processes. As a filter, we selected genes causing the enrichment of more than one process in at least half of the mice.

Interestingly, we observed that *Ppargc1a* (Peroxisome proliferator-activated receptor gamma, coactivator 1 alpha), a transcriptional coactivator, was frequently involved in the up-regulation of energy and lipid metabolism in long-lived mice **(Figure 3B, red labels)**. Given that cells ectopically expressing *Ppargc1a* display resistance to oxidative stress (St-Pierre et al., 2006; Valle et al., 2005), activation of *Ppargc1a* in long-lived mice may explain why these mice maintain a high activity of the electron transport chain without causing oxidative damage.

Also involved in the up-regulation of energy and lipid metabolism we found *Mif* (Macrophage migration inhibitory factor), a cytokine whose increased expression has been noticed not only in long-lived dwarf mice but also in caloric and methionine restricted mice (Miller et al. 2002; Miller et al. 2005). In short-lived mice, *Aldh5a1* (aldehyde dehydrogenase family 5) and *Idh1* (isocitrate dehydrogenase 1) were down-regulated and involved in several processes associated with energy and lipid metabolism **(Figure 3C, red labels)**. Consistently, mice with mutations in these genes display premature death and increased oxidative stress (Hogema et al., 2001; Itsumi et al., 2015; Latini et al., 2007).

We further compared the genes leading the regulation of energy and lipid metabolism in long- and short-lived mice **(Figure 3B-C)** and we identified 5 genes in common: cytochrome c oxidase subunit 5A (*Cox5a*), cytochrome c-1 (*Cyc1*), NADH:ubiquinone oxidoreductase subunit B10 (*Ndufb10*), NADH:ubiquinone oxidoreductase core subunit V2 (*Ndufv2*) and ubiquinol-cytochrome c reductase (*Uqcrfs1*). We directly compared their normalised fold change in long- and short-lived mice using an unpaired two-sample Wilcoxon test. The expression of all 5 genes was regulated in opposite directions between both groups of mice and the differences were statistically significant (p < 0.05) **(Supplementary Figure 3)**.

In short-lived mice, most of the differentially expressed processes were down-regulated and linked with the mitochondria, which may be indicative of mtDNA damage. To probe this idea, we analysed the gene sets enriched in polymerase γ mutant mice (*Polg^D257A/D257A^*), which display a 2500-fold increase in mtDNA mutations compared to wild-type mice (Khrapko and Vijg, 2007). Indeed, we observed that in muscle and brain, the gene sets in Figure 2A were strongly down-regulated, showing that short-lived mice induce a transcriptomic signature matching that mutant mice with exacerbated mtDNA mutations **(Supplementary Figure 4)**.

We also identified gene sets changing expression specifically in long- or short-lived mice. In long-lived mice, consistent with the up-regulation of the electron transport chain, we also observed an up-regulation of processes linked with ATP synthesis **(Supplementary Figure 5)**. This result is in line with a previous study reporting an increase in ATPase activity in long-lived mice (Choksi et al., 2011). We also observed up-regulation of expression of genes involved in thermogenesis, another process activated by *Ppargc1a* in response to cold exposure (Gill and La Merrill, 2017) (Puigserver et al., 1998). Thus, *Ppargc1a* may be activated by the reduced core body temperature typical of long-lived dwarf mice due to a higher body surface to body mass ratio (Ferguson et al., 2007; Hauck et al., 2001; Hunter et al., 1999). Another well-recognised stimulator of oxidative metabolism and ATP production is calcium (Glancy et al., 2013; Griffiths and Rutter, 2009; Tarasov et al., 2012). Consistently, we observed an up-regulation of several genes involved in calcium homeostasis. Unfortunately, there is currently no evidence of the effect on longevity of calcium treatment in mammals. Among the down-regulated gene sets in long-lived mice, we observed several linked with the response to endoplasmic reticulum (ER) stress, including the unfolded protein response (UPR) **(Supplementary Figure 6)**. This down-regulation may reflect lower levels of ER stress, as it has been observed in long-lived *daf-2(e1370)* worms and under caloric restriction (Henis-Korenblit et al., 2010; Matai et al., 2019). In short-lived mice, we only identified two additional groups of gene sets down-regulated and they were related to nucleic acid metabolism and biosynthesis **(Supplementary Figure 7)**. In line with this observation, previous studies have demonstrated a correlation between mitochondrial dysfunction and aberrant biosynthesis of nucleotides (Desler et al., 2007).

### 2.3. Interventions that shorten lifespan resemble the ageing transcriptome

We next compared the transcript profiles of the long-and short-lived mutant mice with profiles characteristic of normal ageing in multiple tissues. We performed functional enrichment analysis using age-related genes from wild-type mice **(see Methods 4.2)**. Changes during normal ageing resembled mostly the transcriptomes of the short-lived mouse models **(Figure 3A, ‘Ageing’ column)**. To test if these similarities also existed at the gene level, we calculated the transcriptome-wide correlations between each mouse model of ageing and the ageing transcriptome. On average, we observed a positive and statistically significant correlation between the transcriptomes of short-lived mice and the changes during ageing (**Figure 4, left panel)**, while interventions that lengthened life displayed a correlation close to zero (mean *r_s_* = 0.005). We further analysed the correlations at the pathway level using enrichment scores and obtained a similar result **(Figure 4, right panel)**. Overall, this analysis supports the hypothesis that accelerated ageing models reproduced partially the molecular changes observed during normal ageing.

**Figure 4.**
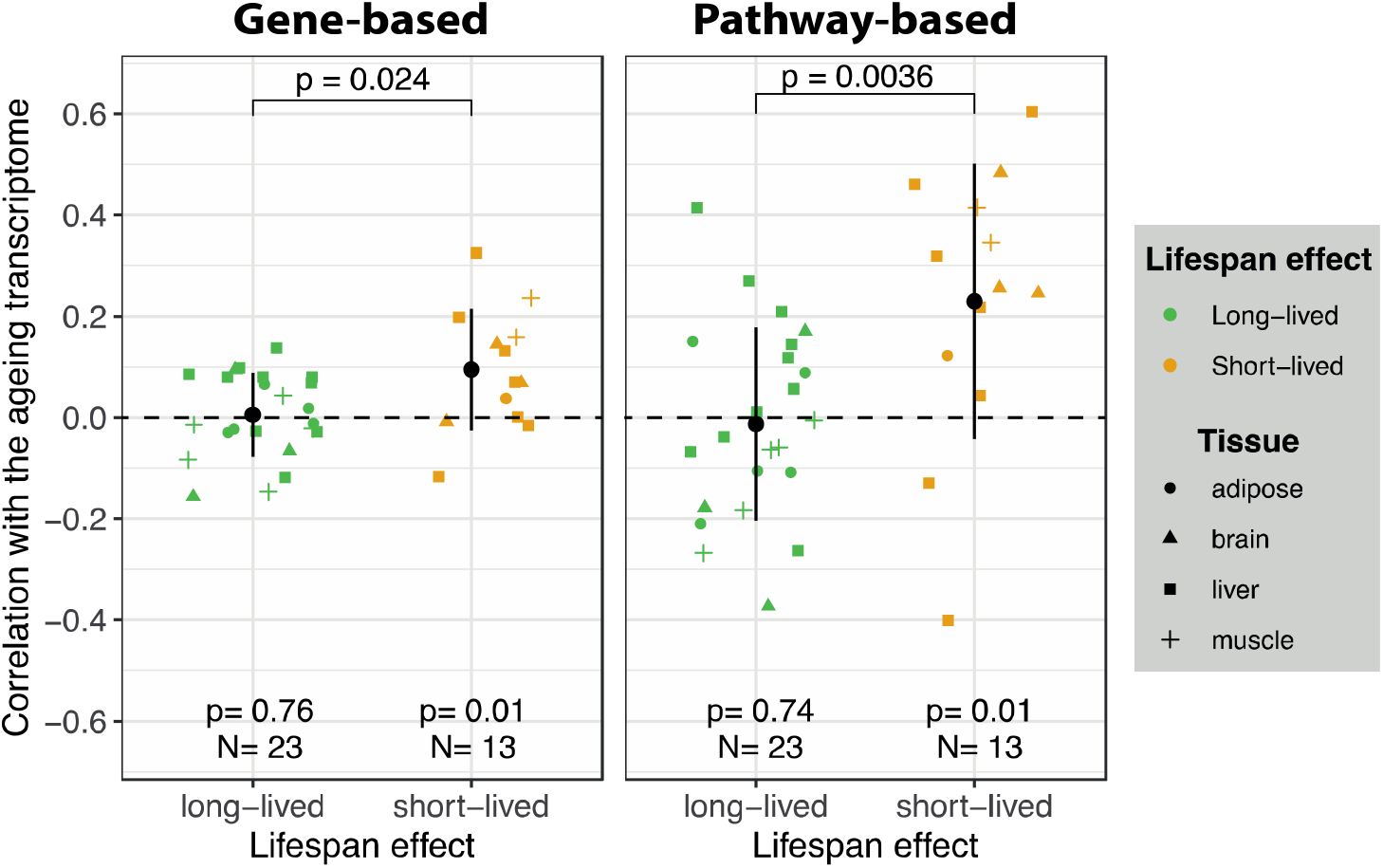
Gene and pathway-based correlations between the transcriptome of ageing-related interventions and that induced by ageing. Each point represents one intervention, and the shapes indicate the tissue from which the transcriptome was derived. Transcriptomic changes in the mouse models of ageing were compared with the changes during ageing on the same tissue. Error bars show one standard deviation from the mean. P-values below were calculated using a t-test with a population mean equal to zero as the null hypothesis. P-values at the top are for the difference between the groups using an unpaired, two-samples Wilcoxon-test.

### 2.4. Identification of potential novel genetic interventions affecting lifespan

We next investigated the possible use of gene sets consistently associated with lifespan **(Figure 3A)** to identify other genetic interventions that could affect ageing. We identified publicly available datasets for other mouse mutants and examined their correlation with the transcriptomes of long-lived mice, short-lived mice and normal ageing **(Figure 5)**. We predict that genetic interventions more positively correlated with long-lived mice will lengthen lifespan, whereas mutants more strongly correlated with short-lived mice and ageing will shorten lifespan. From the 51 gene mutants analysed, 23 showed a higher correlation with long-lived mice and 28 a more positive correlation with short-lived mice. Among the 27 mutants with positive correlation with the ageing transcriptome, 21 (77%) were positively correlated short-lived mice, and 18 (66%) were negatively correlated with long-lived mice. Similarly, from the 24 mutants negatively correlated with ageing, 14 (58%) showed the same trends in short-lived mice, and 21 (87%) displayed the opposite trend in long-lived mice.

**Figure 5.**
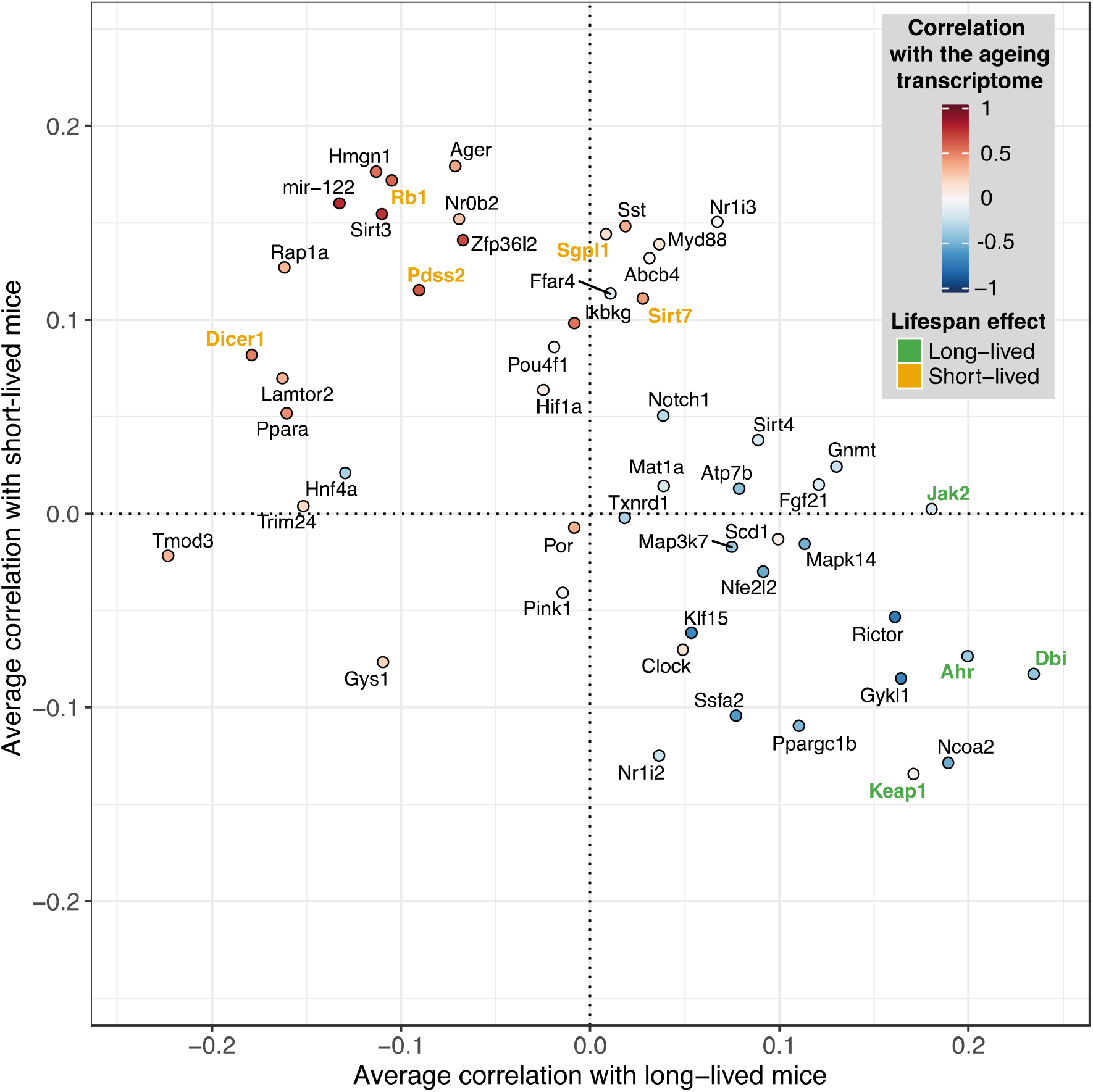
Correlation between the transcriptomes of long- and short-lived mice and that induced by 51 different gene knockouts. Correlations were calculated using enrichments scores of the gene sets in Figure 3A. The colours of the dots represent the correlations with the ageing transcriptome. Labels represent the name of the gene knocked out. Gene knockouts known to lengthen or shorten lifespan in animal models are coloured in green and yellow, respectively.

To determine if the correlational approach we used to identify potential genetic interventions into ageing had external validity, we investigated whether the mutants were previously associated with changes in lifespan in the literature and the GenAge database (De Magalhães and Toussaint, 2004). We also determined if mutations in the orthologues of these genes in *C. elegans* and *Drosophila melanogaster* showed effects on lifespan. Among the 51 gene mutants analysed, we found experimental evidence matching our predictions in 9 cases (Fisher’s exact test p = 0.017). All genetic interventions with experimental evidence to lengthen lifespan showed a more strongly positive correlation of changes in gene expression with those seen in long-lived mice. For instance, the transcriptome of *Jak2* knockout mice showed an average *r_s_* = 0.18 with long-lived mice, but a correlation close to zero against short-lived mice (*r_s_* = 0.002). Consistently, fruit flies with a loss of function mutations in the *hop* gene (orthologue of *Jak2*) live on average 17% longer than wild-type flies (Larson et al., 2012). Similarly, transcriptomic changes induced by *Keap1* knockout in mice were positively correlated with long-lived mice (*r_s_* = 0.17) and negatively correlated with short-lived mice (*r_s_* = −0.13). As expected, *keap1* loss of function mutations extends lifespan of fruit flies by 8-10% (Sykiotis and Bohmann, 2008). The transcriptome of *Ahr* and *Dbi* knockout mice also displayed a positive correlation with long-lived mice and negative correlation with short-lived mice and ageing. Confirming our predictions, *C. elegans* carrying a loss of function allele of *ahr-1* (*Ahr* in mice), display extended lifespan and increase motility and stress resistance (Eckers et al., 2016). Similarly in worms, knockdown of either *acpb-1* or *acbp-3* (*Dbi* in mice), extends lifespan (Shamalnasab et al., 2017). Among the interventions with a more positive correlation with short-lived mice and ageing, we found evidence of premature death in 6 cases, including *Sirt7*, *Dicer1*, *Pdss2*, *Rb1* and *Sgpl1* knockout mice (Frezzetti et al., 2011; Lin et al., 2011; Lyon and Hulse, 1971; Schmahl et al., 2007; Vakhrusheva et al., 2008). The negative effects on lifespan of *Dicer1* and *Sgpl1* have been also shown in *C. elegans*, as loss of *dcr-1* (*Dicer1* in mice) shorten maximum lifespan by 20% (Mori et al., 2012) and RNAi knockdown of *spl-1* (*Sgpl1* in mice), reduces median lifespan by 22% (Samuelson et al., 2007). Overall, the experimental evidence matching with our predictions support the use of gene sets describing mitochondrial metabolism to predict the effects of genetic interventions on lifespan.

## 3. Conclusion

In this study, we collected and analysed publicly available microarray and RNA-seq data of 18 interventions that affect the ageing process and cause changes in lifespan, together with transcript profiles of normal ageing. The transcriptomes were more similar between interventions with the equivalent effects on lifespan, especially if they targeted the same pathway or process. We also detected positive, but weaker, correlations between interventions with opposite effects on lifespan, in line with previous studies (Schumacher et al., 2008). The biggest correlation found in this case, was an undiscovered one between *Lmna^G609G/G609G^* and *Rps6kb1^−/−^*, in the liver and adipose tissue. Interventions like *Akt1^+/−^* (long-lived), *Myc^+/−^* (long-lived) and *Terc^−/−^* (short-lived) did not produce changes in gene expression that correlated with those from other interventions, suggesting the existence of different mechanisms to lengthen and shorten lifespan.

Based on functional enrichment analysis, we identified 58 gene sets (i.e. GO terms), which behaved consistently and showed opposite changes in gene expression in long- and short-lived mouse models. The data implicated mitochondrial metabolism as a key determinant of lifespan. As in short-lived mice, we also detected a transcriptomic down-regulation of mitochondrial metabolism with age in wild-type mice, confirming its relevance for normal ageing and supporting the hypothesis that models of accelerated ageing approximate normal ageing at the molecular level.

Finally, comparing the gene sets associated with lifespan and ageing with those from mouse mutants with no known association with ageing, we found several gene knockouts that were positively correlated with expression changes in long-lived mice and showed consistent experimental evidence from the literature. Our predictions will therefore inform future work on the ageing of these mutant mice.

### 3.1 Limitations and future perspectives

An inherent limitation of the study is that is biased towards interventions in the nutrient-sensing pathways or the DNA repair system, which have been most extensively studied. However, this allowed us to establish a clear relationship between ‘genomic instability and deregulated nutrient sensing’ and ‘mitochondrial dysfunction’, another hallmark of ageing. To develop a complete picture of the interconnections between the hallmarks of ageing, transcriptomic data from interventions targeting other hallmarks of ageing are required.

Also, most of our observations are extracted from the liver transcriptome because it was the only tissue with enough transcriptome data to identify robust trends across interventions. However, as we reported in Section 2.1, genetic interventions may have tissue-specific effects, meaning that the comparison of transcriptomic profiles is likely to be useful only when is done between the same tissue or cell type. Thus, further research in other tissues is necessary to determine the level of conservation of the mechanisms we found associated with lifespan.

## 4. Methods

### 4.1. Dataset collection

We obtained a list of short-lived and long-lived mouse models of ageing from Folgueras et al. (2018) (Folgueras et al., 2018). Then, using the gene symbol of each mutant we searched in the meta-databases *OmicsDI* (Perez-Riverol et al., 2017) and *All of gene expression* (Bono, 2020) for microarray and RNA-seq datasets from mice. Using the *Mouse Genome Database* (Bult et al., 2019), we only included studies with raw data available where the genotype of the wild-type matched the genotype of the mouse model of ageing and where multiple samples per condition were available. We focused on datasets from liver, adipose, muscle and brain as these were the most common tissues among the datasets we found. The samples in each study were grouped into datasets with the same sex, age and tissue, resulting in 57 datasets from 26 studies that were processed separately **(Supplementary Table 1)**. To investigate the gene expression changes during ageing, we used the dataset with the largest sample size for each tissue **(Supplementary Table 5)**.

### 4.2. Differential expression analysis

For microarrays, we obtained the raw data from the *Gene Expression Omnibus* (GEO) and *Array Express* database using the R packages GEOquery (Davis and Meltzer, 2007) and ArrayExpress (Kauffmann et al., 2009), respectively. Affymetrix array data were processed using the RMA algorithm from the package oligo (Carvalho and Irizarry, 2010) to perform background correction, quantile normalisation and summarization. We carry out the summarization using the core genes for exon and gene arrays. For Illumina and Agilent microarrays, we employed background correction and quantile normalisation using the limma package (Ritchie et al., 2015). We removed probes mapping to multiple genes and kept the probe with the highest average expression across samples if multiple probes mapped to one gene. We then fitted a linear model of gene expression versus genotype for each gene and calculated the summary statistics using empirical Bayes. P-values were adjusted for multiple testing using the Benjamini-Hochberg method (Benjamini and Hochberg, 1995) to obtain the false discovery rate. We calculated the gene expression changes of age in the same way, with the difference that chronological age was used as the independent variable instead of genotype.

RNA-seq reads were obtained from the *Sequence Read Archive* (Leinonen et al., 2011). We mapped raw reads to the mouse genome sequence (GRCm38.p6) using STAR (Dobin et al., 2013) and counted mapped reads using featureCounts from the package Rsubread (Liao et al., 2019). We removed genes with less than 10 counts on average across samples. Differential expression analysis was then performed using DESeq2 (Love et al., 2014) controlling the false discovery rate using the Benjamini-Hochberg method.

We further performed a principal component analysis on each dataset using the normalised gene expression values **(Supplementary Figure 8)** and did not observe any sample with outlier levels of expression sufficient to warrant exclusion from the analysis. We also examine which experimental variables (i.e. study, genotype, tissue, sex, age, lifespan effect, technology) accounted for the transcriptional differences between the datasets. We performed a principal component analysis of the quantile normalised fold changes of 4074 genes detected across all 57 datasets **(Supplementary Figure 9)** and then tested which variables explained the differences observed in the first two principal components, applying a multivariate analysis of variance (MANOVA). As expected, datasets from the same study, genotype and tissue tended to correlate more than expected by chance **(Supplementary Table 6)**.

To obtain gene expression data from genetic interventions not previously associated with ageing we downloaded the metadata from all single gene perturbation signatures for mice from the CREEDS database (http://amp.pharm.mssm.edu/CREEDS/#downloads) (Wang et al. 2016). We manually removed interventions not coming from mouse liver or already included in our study. The remaining interventions were processed with GEO2R (https://www.ncbi.nlm.nih.gov/geo/geo2r/).

### 4.3. Transcriptome-wide correlation analysis

Using the log2 fold changes from the differential expression analysis, we calculated the correlation between the datasets using Spearman’s rank method (Spearman, 1904) considering genes in common between the pair of datasets. On interventions with multiple datasets, we averaged the correlations first between datasets from the same study and intervention, and then with the same intervention across different studies. Heatmaps visualising correlations were created using ComplexHeatmap (Gu et al., 2016).

### 4.4. Functional enrichment and consistency analysis

We performed functional enrichment analysis of each dataset separately using the function gseGO from the package ClusterProfiler (Yu et al., 2012) on genes ranked by the sign of the log2 fold changes multiplied by the logarithm of the p-values from the differential expression analysis. Based on this rank-ordered list of genes, we further assessed if genes in each gene ontology term were more likely to be up or down-regulated than what is expected by chance based on 10,000 permutations. We calculated an enrichment score for each pathway and each intervention by multiplying the obtained log10(p-value) by the 1 or −1 depending on the direction of the change. To identify GO terms with consistent changes, we calculated for each GO term the median rank of the enrichment score across short-lived or long-lived mice and we compare it against a random distribution of median ranks with the same number of interventions. P-values were obtained by dividing the observed median rank by the total number of random median ranks generated (i.e. 1e^6^). We adjusted the p-values for multiple testing using the Benjamini-Hochberg method.

To visualise gene ontology terms specific to long- or short-lived mice, we calculated the similarity between the consistently varying terms using the overlap coefficient (oc). We constructed a network connecting the terms (nodes) were edges represent pairs of terms with an oc > 0.4. Clusters with less than 5 nodes were excluded.

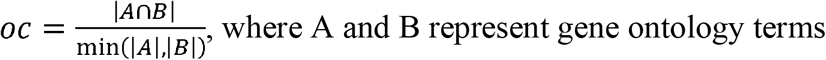

## Funding

This work was funded by Comisión Nacional de Investigación Científica y Tecnológica - Government of Chile (CONICYT scholarship) (MF), EMBL (HMD, DKF, JMT), and the Wellcome Trust [WT098565/Z/12/Z] (LP, JMT).

## Notes

### Competing Interest Statement

The authors have declared no competing interest.

