## Supplementary material for "Transcriptomic profiling of long- and short-lived mutant mice implicates mitochondrial metabolism in ageing and shows signatures of normal ageing in progeroid mice"

Supplementary Tables

**Supplementary Table 1.** Transcriptomic datasets of the mouse models of ageing.

**Supplementary Table 2.** Transcriptomic datasets of genetic interventions not previously associated with ageing. The last two columns represent the sample identifiers of controls and mutant mice.

**Supplementary Table 3.** Gene sets with consistent transcriptomic changes in long-lived mice.

**Supplementary Table 4.** Gene sets with consistent transcriptomic changes in short-lived mice.

**Supplementary Table 5.** Transcriptomic datasets of mice at different ages.

**Supplementary Table 6.** Experimental variables influencing the transcriptomic differences between the datasets

**Supplementary Figures**

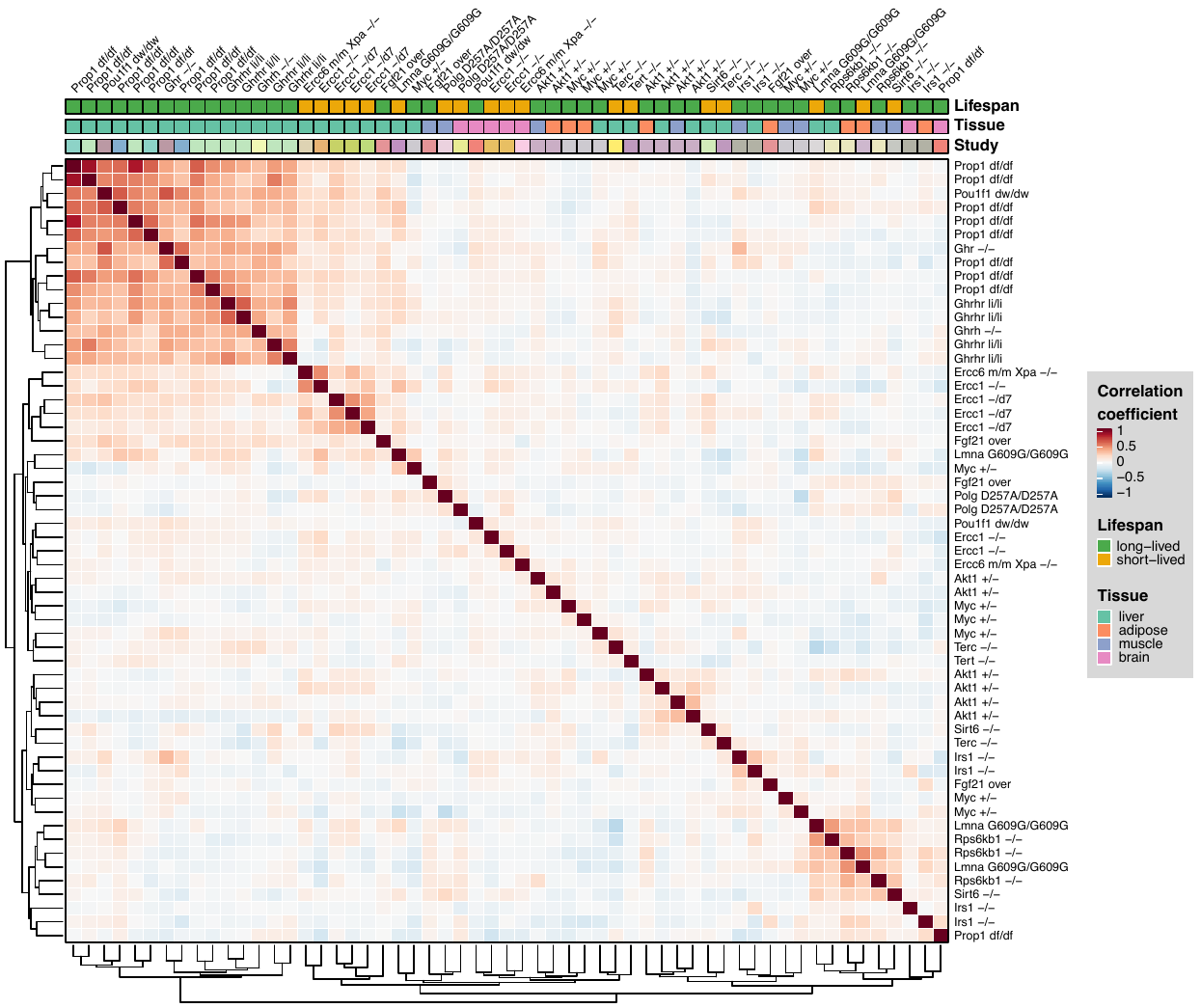

**Supplementary Figure 1.** Spearman’s rank correlation coefficients between the transcriptome of mouse models of ageing.

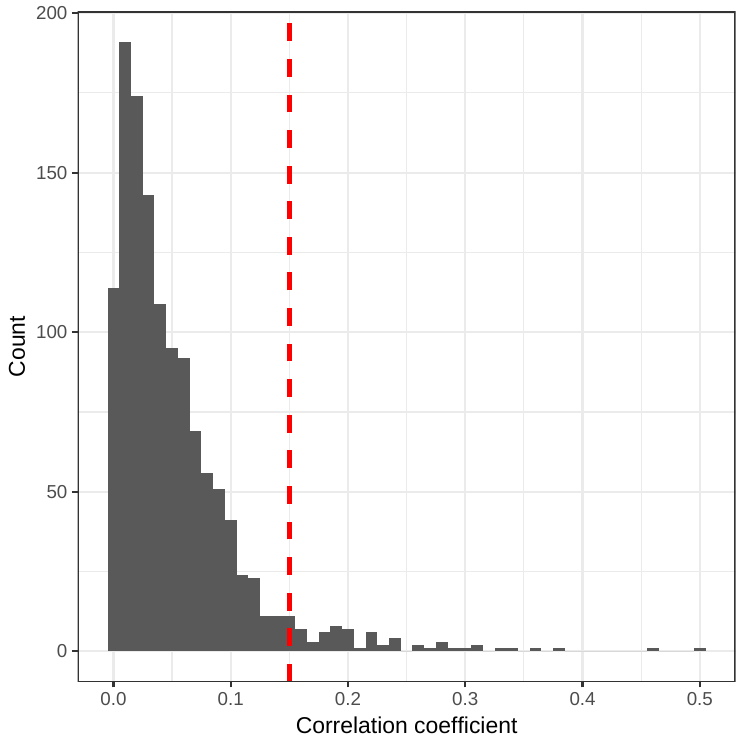

**Supplementary Figure 2.** Distribution of pairwise correlation between the transcriptomes of mutants not previously associated with ageing.

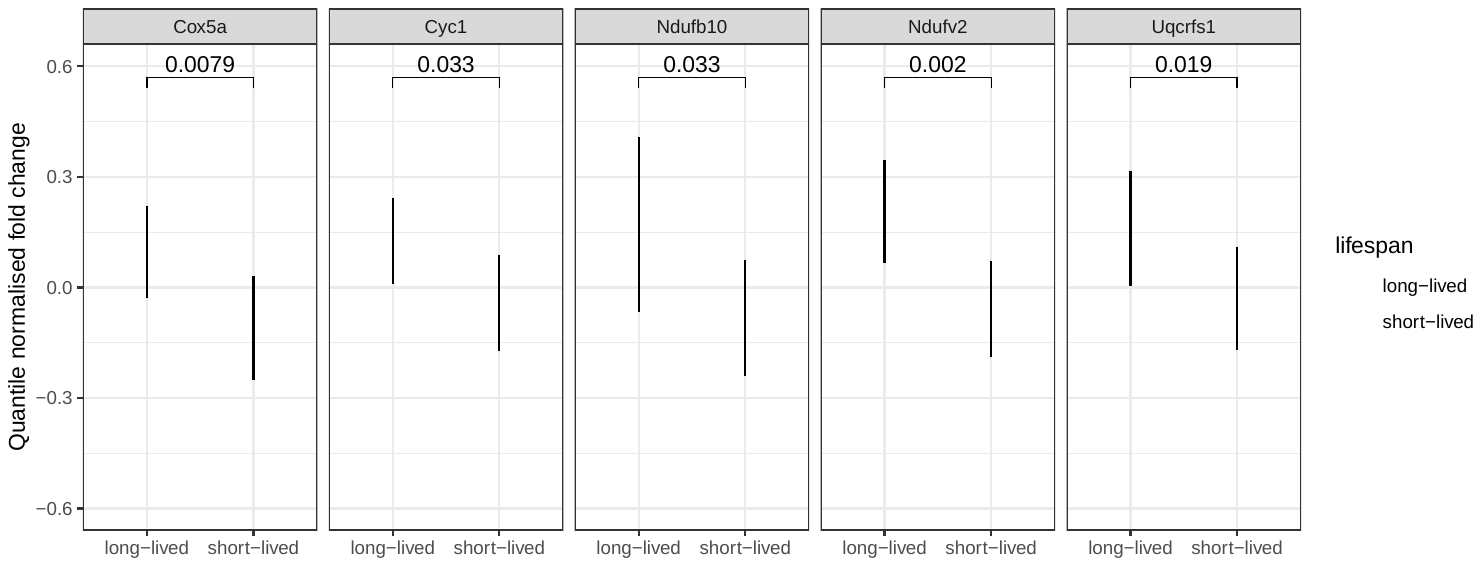

**Supplementary Figure 3.** Expression of genes leading the regulation of energy and lipid metabolism in long- and short-lived mice.

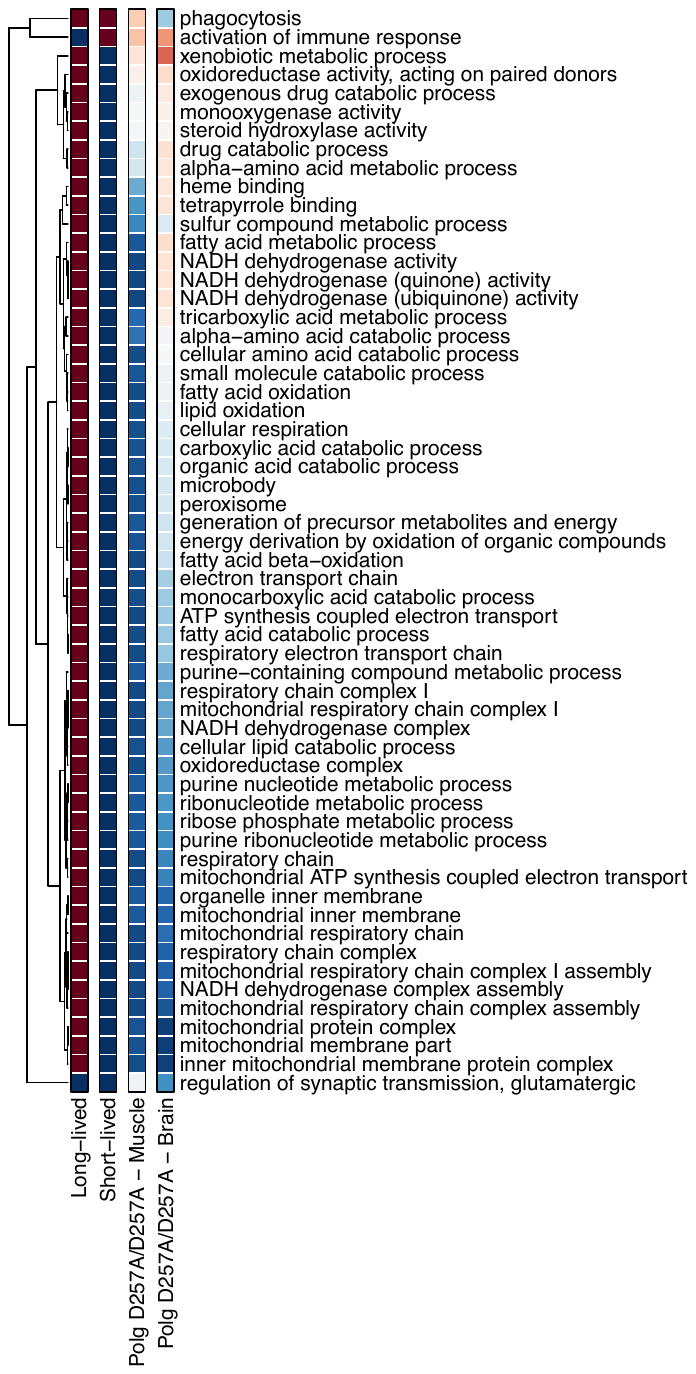

**Supplementary Figure 4.** Transcriptional changes in *Polg^D257A/ D257A^* mutant mice in gene sets associated with lifespan and ageing.

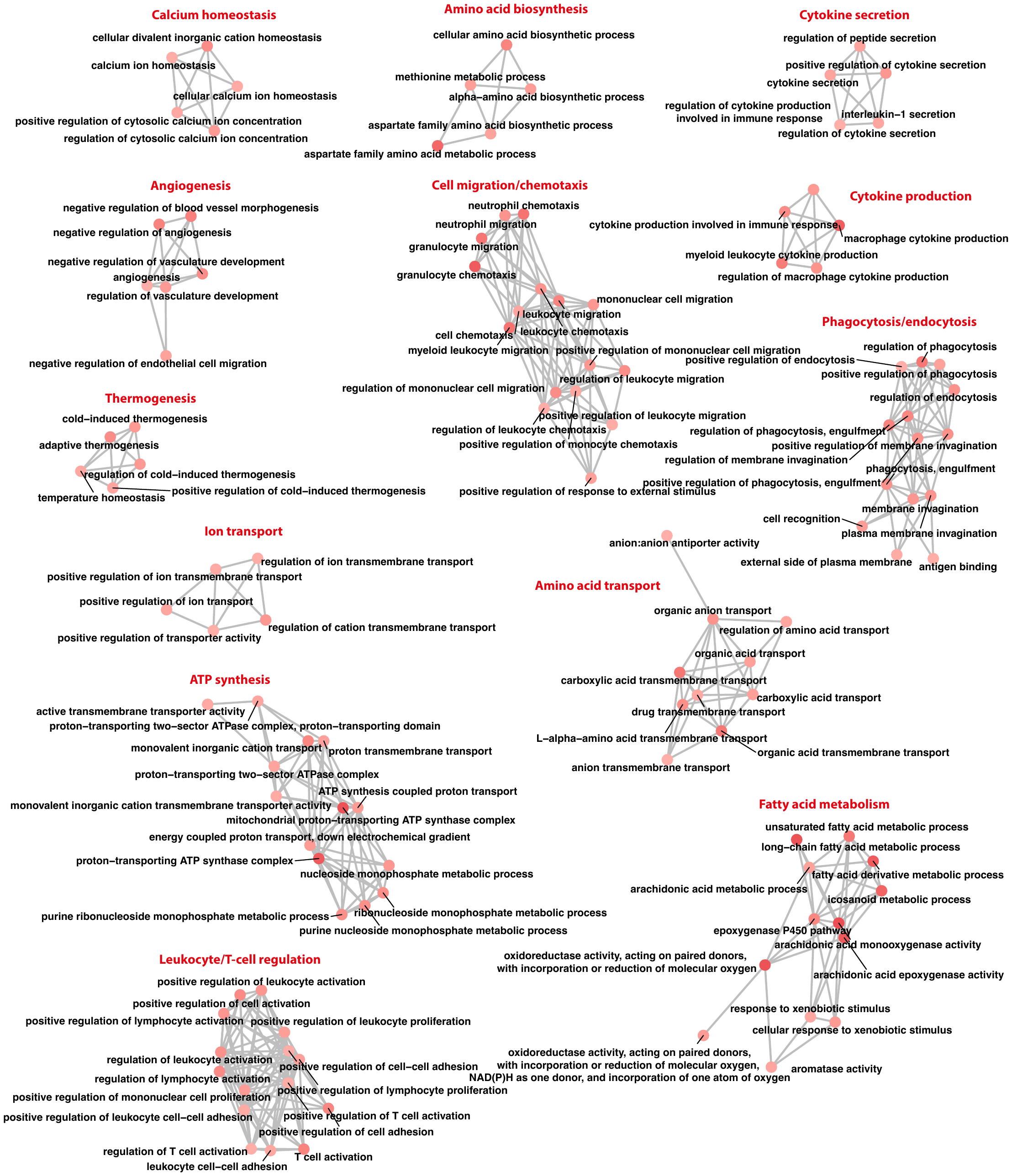

**Supplementary Figure 5.** Gene set up-regulated in long-lived mice. Nodes represent GO terms and edges indicate shared genes between the terms. The colour of the circles denotes the significance of the consistency across mutant mice.

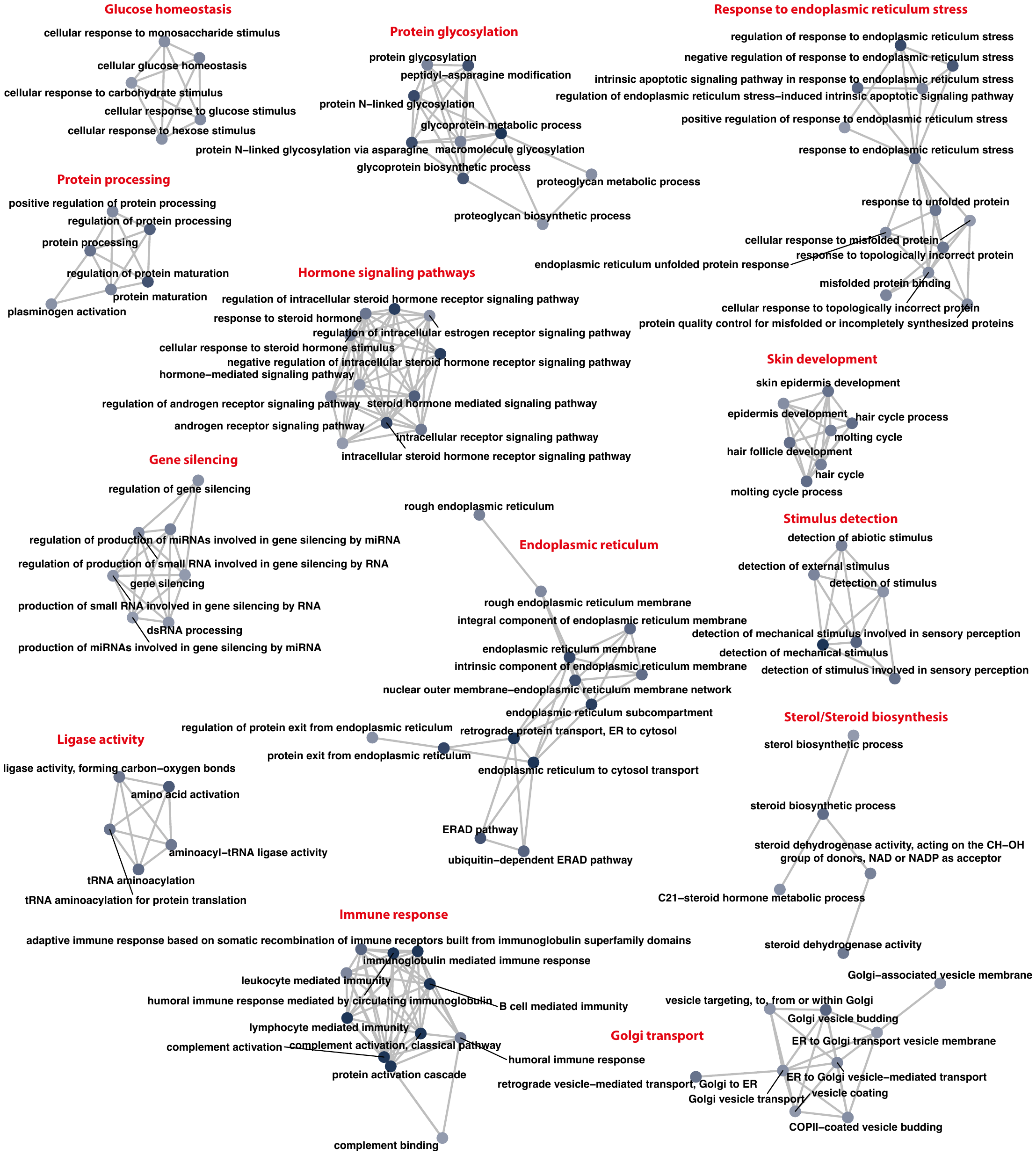

**Supplementary Figure 6.** Gene set down-regulated in long-lived mice. Nodes represent GO terms and edges indicate shared genes between the terms. The colour of the circles denotes the significance of the consistency across mutant mice.

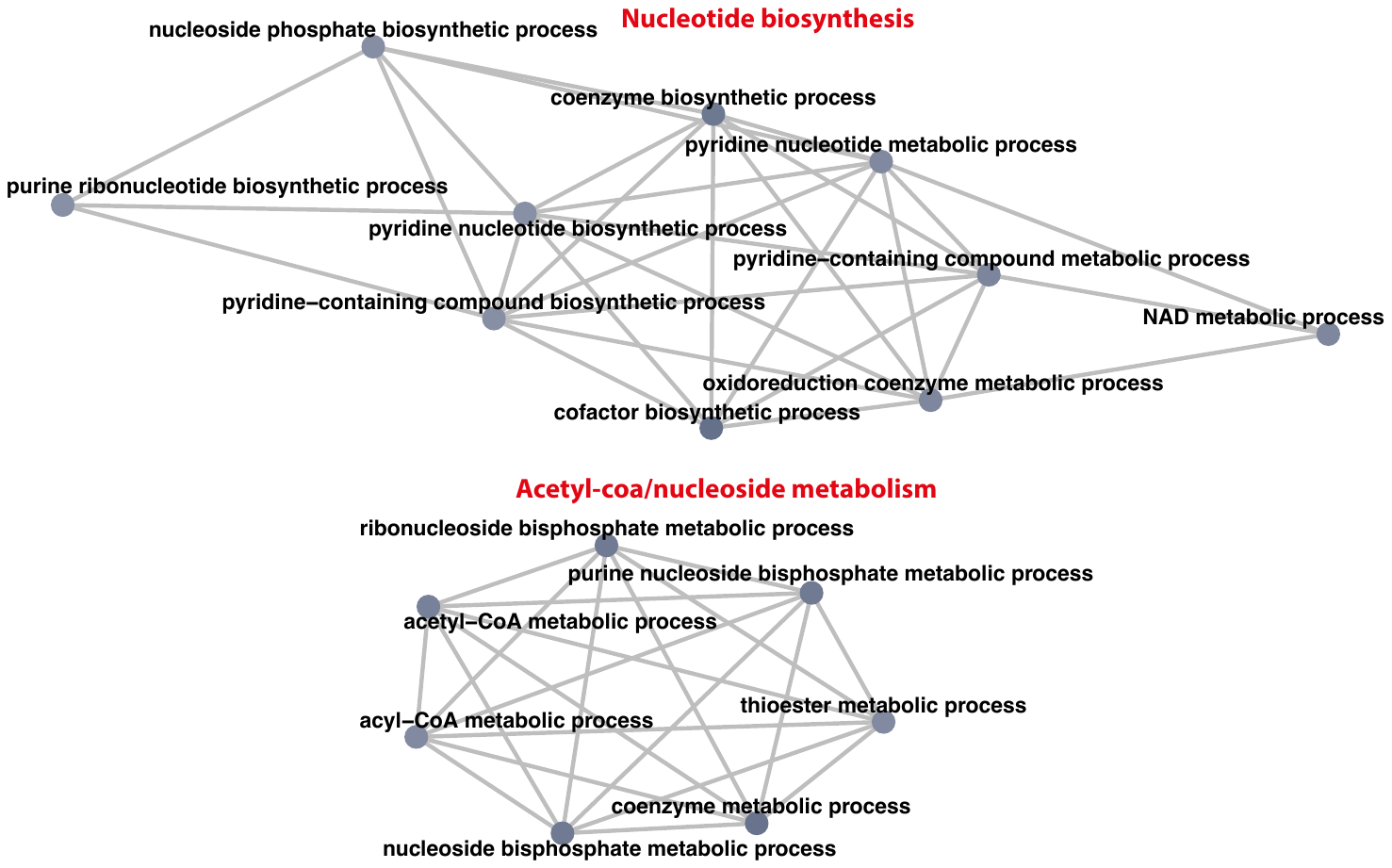

**Supplementary Figure 7.** Gene set down-regulated in short-lived mice. Nodes represent GO terms and edges indicate shared genes between the terms. The colour of the circles denotes the significance of the consistency across mutant mice.

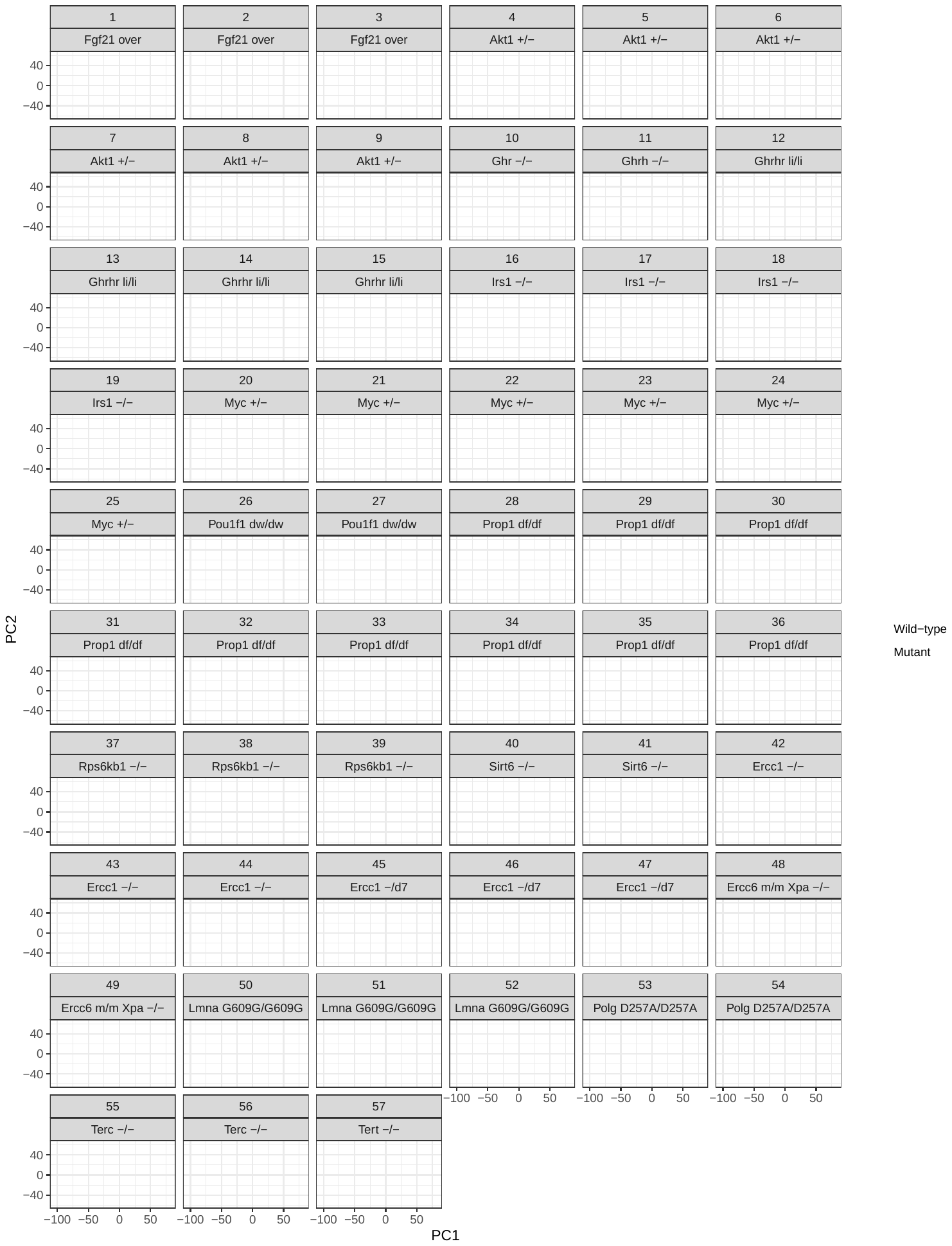

**Supplementary Figure 8.** Principal component analysis of the datasets based on expression levels. The samples are coloured by the genotype.

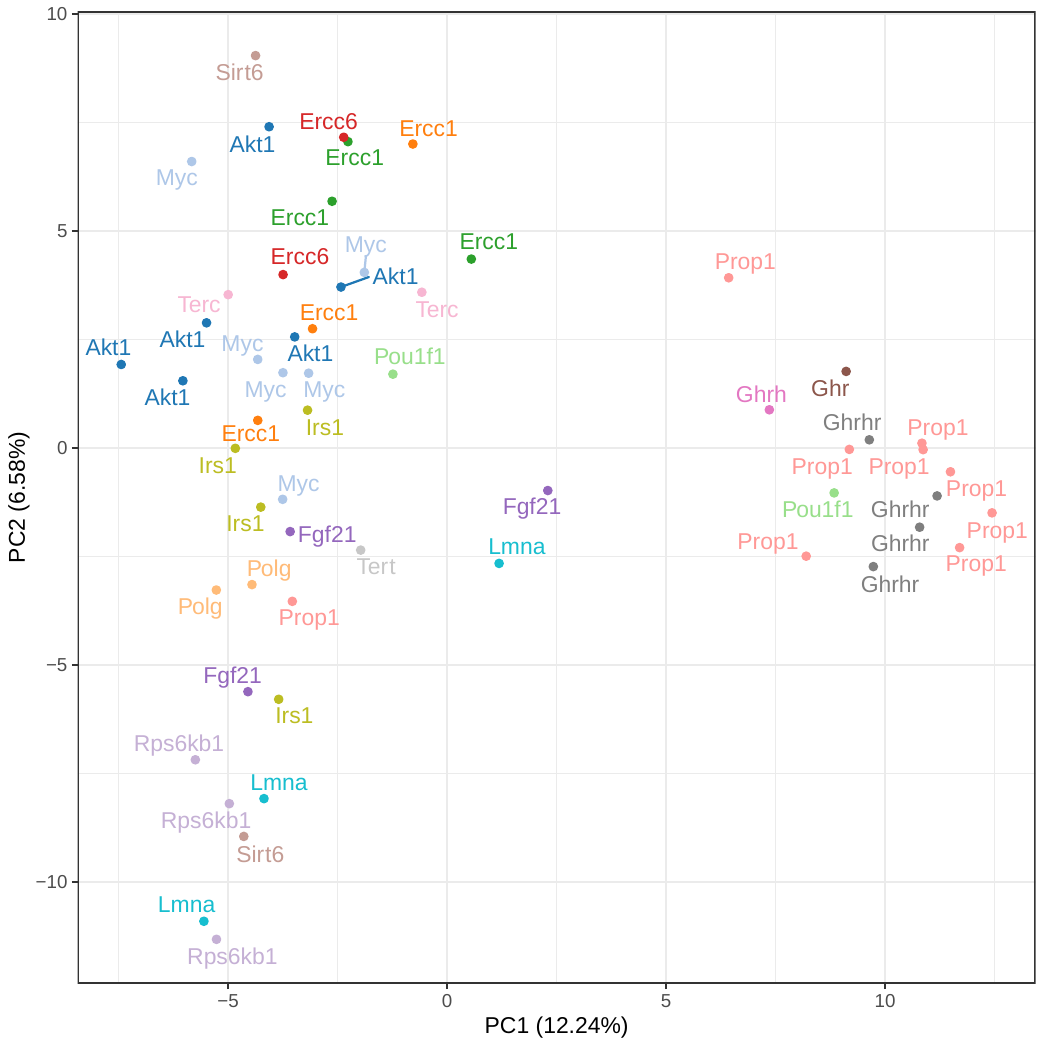

**Supplementary Figure 9.** Principal component analysis of the interventions based on the normalised fold changes from the differential expression analysis. The labels and colour indicate the genes altered. In parenthesis on the axis is the percentage of the variance explained by each principal component.
